# Evolution within a given virulence phenotype (pathotype) is driven by changes in aggressiveness: a case study of French wheat leaf rust populations

**DOI:** 10.1101/2022.08.29.505401

**Authors:** Cécilia Fontyn, Kevin JG Meyer, Anne-Lise Boixel, Ghislain Delestre, Emma Piaget, Corentin Picard, Frédéric Suffert, Thierry C Marcel, Henriette Goyeau

## Abstract

Plant pathogens are constantly evolving and adapting to their environment, including their host. Virulence alleles emerge, and then increase, and sometimes decrease in frequency within pathogen populations in response to the fluctuating selection pressures imposed by the deployment of resistance genes. In some cases, these strong selection pressures cannot fully explain the evolution observed in pathogen populations. A previous study on the French population of *Puccinia triticina*, the causal agent of wheat leaf rust, showed that two major pathotypes — groups of isolates with a particular combination of virulences — predominated but then declined over the 2005-2016 period. The relative dynamics and the domination of these two pathotypes — 166 317 0 and 106 314 0 —, relative to the other pathotypes present in the population at a low frequency although compatible, i.e. virulent on several varieties deployed, could not be explained solely by the frequency of *Lr* genes in the landscape. Within these two pathotypes, we identified two main genotypes that emerged in succession. We assessed three components of aggressiveness — infection efficiency, latency period and sporulation capacity — for 44 isolates representative of the four *P. triticina* pathotype-genotype combinations. We showed, for both pathotypes, that the more recent genotypes were more aggressive than the older ones. Our findings were highly consistent for the various components of aggressiveness for pathotype 166 317 0 grown on Michigan Amber — a ‘naive’ cultivar never grown in the landscape — or on Apache — a ‘neutral’ cultivar, which does not affect the pathotype frequency in the landscape and therefore was postulated to have no or minor selection effect on the population composition. For pathotype 106 314 0, the most recent genotype had a shorter latency period on several of the cultivars most frequently grown in the landscape, but not on ‘neutral’ and ‘naive’ cultivars. We conclude that the quantitative components of aggressiveness can be significant drivers of evolution in pathogen populations. A gain in aggressiveness stopped the decline in frequency of a pathotype, and subsequently allowed an increase in frequency of this pathotype in the pathogen population, providing evidence that adaptation to a changing varietal landscape not only affects virulence but can also lead to changes in aggressiveness.

## INTRODUCTION

Plant diseases and pests cause crop damage accounting for up to 40% of yield losses (Boonekamp, 2012). Pathogenicity, or the ability of plant pathogens, especially fungi, to cause disease, is generally broken down into a qualitative term, ‘virulence’, and a quantitative term, ‘aggressiveness’ (Lannou, 2012). Virulence is defined as the capacity of the pathogen to infect its host (compatible interaction), as opposed to avirulence, which expresses the resistance of the host (incompatible interaction), according to the gene-for-gene model (Flor, 1971). A virulence phenotype, also known as a pathotype or race, is defined by a virulence profile: two pathogenic isolates are considered to belong to the same pathotype if they have the same combination of virulences. Aggressiveness, the quantitative variation of pathogenicity on a compatible host (Pariaud *et al*., 2009a), can be viewed as the detrimental impact of a pathogen on its host, leading to damage to the crop plant and, thus, yield losses (Shaner *et al*., 1992; Pariaud *et al*., 2009a; Lannou, 2012). Aggressiveness also determines the rate at which a given disease intensity is reached. Its assessment is intrinsically complex because it is related to various life-history traits of the pathogen specific to its biology and the nature of the symptoms that it produces. The different components of aggressiveness can be measured by evaluating several of these complementary quantitative traits expressed during the host– pathogen interaction. The most widely assessed aggressiveness components for rust pathogens are infection efficiency, latency period and sporulation capacity (Pariaud *et al*., 2009a; Lannou, 2012; Azzimonti *et al*., 2013). A higher infection efficiency will directly cause more host damage, while latency period and sporulation capacity will also increase the parasitic fitness of the pathogen (Shaner *et al*., 1992) by favoring its transmission before damaging the host . However, the relationship between these aggressiveness components of the pathogen and the reduction in crop yield and biomass remains a theoretical assumption that is rarely verified experimentally (Van Roermund and Spitters, 1990).

Infection efficiency is calculated by determining the proportion of unit of inoculum, i.e. spores, that cause a new infection when deposited on compatible host plant tissues (Sache, 1997). The estimation of this component is complex due to technical issues, particularly the need for great precision in the spore deposition process (Lehman & Shaner, 1997), which involves placing a fixed and known number of spores — ideally one by one — on the leaf.

The latency period is the length of time between “the start of the infection process by a unit of inoculum”, i.e. the deposition of a spore on plant tissues, and “the start of production of infectious units”, i.e. first sporulation (Parlevliet, 1979; Madden *et al*., 2007). In rusts, this component is often defined as the length of time between inoculation and the appearance of 50% of the sporulating structures, also known as ‘uredinia’ (Parlevliet, 1975; Johnson, 1980; Pfender, 2001). Latency period estimations therefore require counts of uredinia on at least a daily basis. The latency period is highly temperature-dependent, and its expression in thermal time is therefore recommended, to allow comparisons between trials (Lovell *et al*., 2004).

Sporulation capacity is assessed as the number of spores produced per individual sporulating structure and per unit time (Sache, 1997; Pariaud *et al*., 2009a). Spores can be collected and counted directly (e.g. with a cell counter) or indirectly (e.g. weighed) (Imhoff, 1982; Robert *et al*., 2004; Delmotte *et al*., 2014). However, in rusts, uredinium density affects the number of spores produced (Robert *et al*., 2004) and must, therefore, be taken into account in some analysis (Lannou & Soubeyrand, 2017). Moreover, sporulation is a continuous process, so sporulation capacity is time-dependent. Thus, this component, like several others, is dependent on latency period. The interdependence of traits can be reduced by measuring sporulation capacity at a normalized time point.

Leaf rust caused by *Puccinia triticina* is one of the most damaging wheat diseases, causing high yield losses worldwide (Huerta-Espino *et al*., 2011; Savary *et al*., 2019). Qualitative resistance is the easiest and most effective means of limiting leaf rust epidemics. Eighty-two *Lr* genes have been identified in wheat cultivars, most displaying qualitative interactions (Bariana *et al*., 2022). It has been shown that the deployment of qualitative resistance genes in the landscape exerts a strong selective pressure, acting as an important driver of evolution in *P. triticina* populations (Goyeau *et al*., 2006). This effect was highlighted by surveys of virulence phenotypes (pathotypes), which showed that the corresponding virulence can emerge rapidly after the introduction of a new *Lr* gene into cultivars. For example, in France, virulence against *Lr*28 appeared within only two years of the release of cultivars carrying *Lr28* (Fontyn *et al*., 2022). Adaptation to qualitative resistances occurs rapidly, despite the clonality of the population, through ‘boom-and-bust’ cycles of resistance (McDonald & Linde 2002).

However, qualitative resistance alone cannot fully explain the evolution of the composition of the pathogen population. The occurrence of selection for greater aggressiveness has already been established for various pathogens (Delmas *et al*., 2016; Frézal *et al*., 2018).Milus *et al*. (2009) showed that the replacement of an ‘old’ *Puccinia striiformis* f. sp. *tritici* population by a ‘new’ population could be explained by the greater aggressiveness of isolates from this new population, in addition to a change in its composition in pathotypes. A survey of the French *P. triticina* population from 1999-2002 (Goyeau *et al*., 2006) revealed the domination of a single pathotype (073 100 0), coinciding with a period in which the cultivar landscape was dominated by the cultivar Soissons. Pathotype 073 100 0 was found to be more aggressive on this cultivar than other virulent pathotypes present in the *P. triticina* population during this period (Pariaud *et al*., 2009b). Fontyn *et al*. (2022) recently showed that the domination of the French landscape by two pathotypes — 106 314 0 and 166 317 0 — during the 2005-2016 period could not be fully attributed to the deployment of *Lr* genes. Indeed, several other compatible pathotypes virulent against the *Lr* genes carried by the most widely grown cultivars were present in the landscape, but never reached substantial frequencies. The authors suggested that aggressiveness might drive the evolution of *P. triticina* populations, modifying pathotype frequencies at large spatiotemporal scales. The variation in aggressiveness over time, during the complete lifespan of a pathotype, including its emergence, domination and replacement, has never been investigated.

The objective of this study was to determine if an evolution of aggressiveness can occur within a pathotype of *P. triticina* over a large temporal scale and whether information on aggressiveness allows us to explain retrospectively or even predict changes in pathotypes frequency in the landscape. To this end, we focused on two major pathotypes, 166 317 0 and 106 314 0, identified as good experimental case studies for this purpose, because of their long lifespan and high frequency in the French landscape over the 2005-2016 period. We first characterized isolates of these two pathotypes using microsatellite markers, to identify their genotypic diversity. Within each of the two pathotypes, we then compared the aggressiveness components of isolates from the most frequent genotypes (i) on ‘neutral’ cultivars, i.e. cultivars with no apparent effect on the frequencies of these pathotypes, and, for pathotype 106 314 0 only, (ii) on five cultivars widely grown in France over the study period.

## MATERIALS AND METHODS

### Selection and purification of isolates

Annual surveys of *P. triticina* populations were carried out at INRAE BIOGER over the last two decades. These surveys involve the collection of leaf samples from field (micro)plots sown with a single variety in a network of field trials and nurseries throughout the wheat-growing areas of France (Goyeau *et al*., 2006; Fontyn *et al*., 2022). In total, 2796 leaves were collected from the 10 most cultivated varieties during the 2006-2016 period in the framework of the national survey. Urediniospores were bulk-harvested from each leaf and stored at −80°C. One single-uredinium isolate was selected from each bulk and its pathotype was determined as described by Goyeau *et al*. (2006). In total, 932 isolates were identified as pathotype 106 314 0 and 473 isolates were identified as pathotype 166 317 0 during the national survey. For the purposes of this study, we selected 286 urediniospore bulks of pathotype 106 314 0 collected between 2006 and 2016, and 115 from pathotype 166 317 0 collected between 2013 and 2016 (Table S1 and S2). No bulk were selected for pathotype 166 317 0 before 2013 because the frequency of this pathotype in the landscape was too low. The bulks were defrosted and repurified by the re-inoculation of seven-day-old cv. Michigan Amber wheat seedlings, to obtain 401 new single-uredinium isolates. Before inoculation, the plants were grown in cabinets with air filters in a greenhouse at temperatures between 15 and 20°C, under a 14-h photoperiod (daylight supplemented with 400 W sodium lamps). Plants were treated with 15 mL maleic hydrazide solution (0.25 g maleic hydrazide per liter of H_2_O) to prevent the emergence of secondary leaves and to increase spore production. Inoculated seedlings were placed in a dew chamber at 15°C for 24 h and were then transferred to the greenhouse. One week after inoculation, the seedlings were trimmed such that only one plant with one uredinium remained in each pot. Before sporulation, cellophane bags were placed over the pots to prevent contamination between isolates. Ten days after inoculation, 401 leaf segments, each carrying only one uredinium, were collected for DNA extraction and genotyping with microsatellite markers. Other single-uredinium isolates were also selected from 28 and 16 bulks initially identified as pathotypes 106 314 0 and 166 317 0, respectively. Each of these 44 isolates was pathotyped (Goyeau *et al*., 2006) and, after confirmation that their virulence phenotypes were as expected, genotyped with microsatellite markers. Spores from these 44 isolates were stored at −80°C for further assessments of aggressiveness (Table S3). The various stages in the purification, selection, pathotyping and genotyping of *P. triticina* isolates for which aggressiveness components were evaluated are summarized in Figure 1.

**Figure 1.**
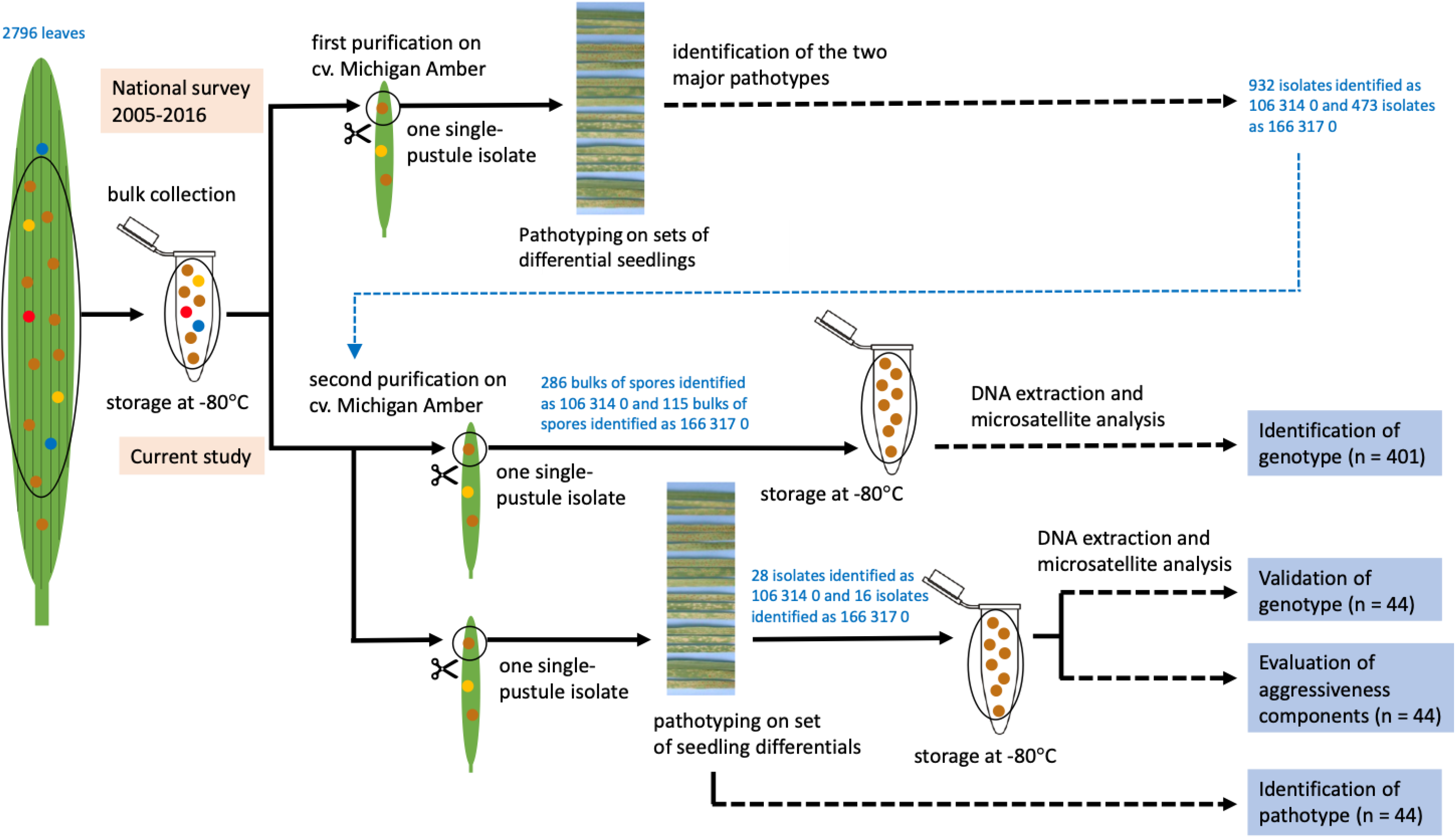
Overview of the purification, selection, pathotyping, and genotyping steps for the *P. triticina* isolates for which aggressiveness components were evaluated.

### Genotyping of isolates with microsatellite markers

#### DNA extraction

DNA was extracted from all the purified isolates in 96-well extraction plates, with Qiagen DNeasy® Plant Mini Kit buffers (Qiagen, Hilden, Germany). To this end, each leaf segment carrying a single uredinium was placed in a Qiagen collection microtube with a tungsten bead and 100 μL of hot AP1 buffer (65°C). Leaf segments were ground by shaking the microtubes in a Retsch® MM400 ball mill twice, for 30 seconds each, at 25 Hz. The tubes were then centrifuged for 1 minute at 3000 x *g*. AP1 buffer supplemented with RNase A and Reagent DX was then added to each tube (300 μL). After mixing, we added 130 μL P3 buffer to the tube, which was then incubated for 10 minutes at −20°C and centrifuged at 4°C for 10 minutes at 20 000 x *g*. The supernatant (200 μL) was transferred to a new tube, to which we added 1 volume of sodium acetate (3 M pH 5) and 3 volumes of isopropanol (100%). After mixing, the tubes were placed at −20°C for 30 minutes and then centrifuged for 20 minutes at 6000 x *g*. The supernatant was removed and 3 volumes of 70% ethanol were added. The tube was placed at −20°C for 5 minutes and was then centrifuged at 4°C for 15 minutes at 6000 x *g*. The pellet containing the DNA was allowed to dry overnight, and was then resuspended in 100 μL ultra-purified water. We transferred 20 µL of the resulting suspension to Qiagen elution microtubes RS in a 96-tube rack, which was sent to Eurofins (Eurofins, Luxembourg) for genotyping. The DNA suspensions concentrations were between 1 and 25 ng/μL.

#### Microsatellite genotyping and analyses

The 401 and 44 single-uredinium isolates were genotyped for 19 microsatellite markers: RB8, RB11, RB12, RB17, RB25, RB26, PtSSR13, PtSSR50, PtSSR55, PtSSR61, PtSSR68, PtSSR91, PtSSR92, PtSSR152, PtSSR154, PtSSR158, PtSSR164, PtSSR173, and PtSSR186 (Duan *et al*., 2003; Szabo & Kolmer, 2007). The microsatellite markers were assembled into two multiplexes of 9 and 10 markers and labeled with four fluorochromes (Table S4) to prevent overlaps between markers with the same range of allele sizes. PCR amplification was performed by Eurofins (Eurofins, Luxembourg), with the following amplification program: 95°C for 5 min and 35 cycles of 95°C for 30 s, 58°C for 30 s, 72°C for 30 s, and then 60°C for 30 min. Each reaction contained 5 μL DNA solution and PCR mixture with Taq Type-it (Qiagen®). PCR products were analyzed by capillarity electrophoresis on an ABl3130xl sequencer. The two-multiplex PCR system was tested and validated on eight reference *P. triticina* DNA extracts from isolates obtained from bread and durum wheat. Chromatograms were visually inspected for all markers and for all individuals with Peak Scanner software version 2.0, before the final assignment of SSR alleles. The chromosome position of SSR markers was determined by performing a blastn analysis (with default parameter values) of the sequence of each primer against the two genome sequences of the Australian isolate Pt76 (Duan *et al*., 2022) with the NCBI blast tool (https://blast.ncbi.nlm.nih.gov/Blast.cgi).

The genotyping of the 401 isolates revealed two major genotypes within each pathotype: 106 314 0-G1 and 106 314 0-G2 for pathotype 106 314 0, and 166 317 0-G1 and 166 317 0-G2 for pathotype 166 317 0. These genotypes were taken into account in the design of subsequent experiments.

### Evaluation of aggressiveness components

#### Experimental design

Three components of aggressiveness, infection efficiency, latency period and sporulation capacity, were assessed for 28 isolates of pathotype 106 314 0 and 16 isolates of pathotype 166 317 0 that had been purified, pathotyped and genotyped (Table S3). These aggressiveness components were measured in the greenhouse on seedlings of two wheat varieties: (i) Apache, a commercial French cultivar characterized as ‘neutral’ in a previous study (Fontyn *et al*., 2022), i.e. with no selection effect on the landscape-pathotype pattern, and (ii) Michigan Amber, considered intrinsically ‘naive’ or ‘neutral’, both because it carries no known leaf rust resistance gene and because it has never been cultivated in France and cannot, therefore, have played any significant role in the evolutionary trajectory of *P. triticina* in France.

It was not possible to perform a large single trial at our facilities, so the 44 isolates were characterized in five successive series, according to the same protocol and under the same experimental conditions. In series 1 (performed in March-April 2021) and 2 (August-September 2021), we tested whether the two genotypes within each *P. triticina* pathotype differed in aggressiveness on Apache and Michigan Amber. The isolates used for these two series were collected between 2005 and 2016 on Apache (Table 1). In series 3 (July-august 2020), 4 (April-May 2021) and 5 (May-June 2021), we tested the difference in aggressiveness between the two genotypes within pathotype 106 314 0 (106 314 0-G1 and 106 314 0-G2) on some or all of the wheat cultivars Aubusson, Premio, Michigan Amber, Sankara, Expert and Bermude. Except Michigan Amber, all were among the 35 most frequently grown cultivars in the French landscape during the 2006-2016 period. The isolates used in these three series were collected in 2012 and 2013, on Aubusson, Premio and Apache (Table 2). The isolates used in series 3 were collected on cultivars Aubusson and Premio and were tested on these two cultivars and Michigan Amber. The isolates used in series 4 were collected on cultivar Apache and were tested on Apache and Michigan Amber. The isolates used in series 5 were collected on cultivar Apache and were tested on cultivars Sankara, Expert and Bermude. In all five series, each isolate × cultivar interaction was replicated eight times, on eight wheat seedlings. Three replicates of the experimental design, including the five series, were established.

**Table 1.** Experimental design, with the allocation of isolates to series 1 and 2, for assessments of the aggressiveness of pathotypes 106 314 0 and 166 317 0. Isolates were collected during the 2005-2016 period, from cultivar Apache, and were tested on Apache and Michigan Amber.

| Pathotype | Genotype<sup>a</sup> | Year of sampling | Number of isolates | Series |
| --- | --- | --- | --- | --- |
| 106 314 0 | 106 314 0-G1 | 2005-2006 | 6 | 1 |
|  |  | 2012-2013 | 3 | 1 |
|  |  | 2015-2016 | 0 | 1 |
|  | 106 314 0-G2 | 2005-2006 | 0 | 1 |
|  |  | 2012-2013 | 2 | 1 |
|  |  | 2015-2016 | 6 | 1 |
| 166 317 0 | 166 317 0-G1 | 2012 | 6 | 2 |
|  |  | 2014 | 6 | 2 |
|  |  | 2016 | 0 | 2 |
|  | 166 317 0-G2 | 2012 | 0 | 2 |
|  |  | 2014 | 0 | 2 |
|  |  | 2016 | 4 | 2 |

<sup>a</sup> Genotyping with 19 SSRs specific to leaf rust (see Table 3)

**Table 2.**
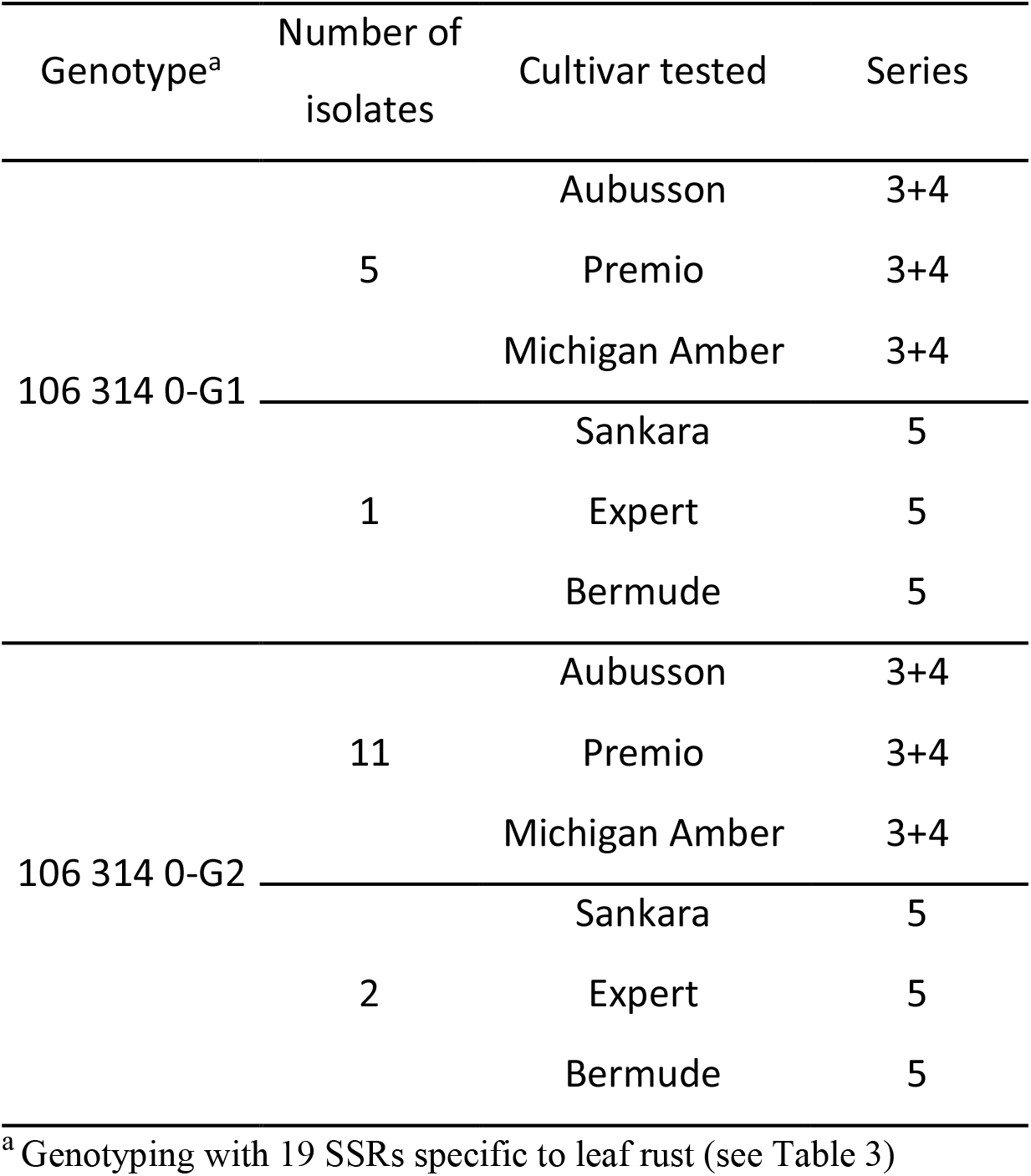
Experimental design, with the allocation of isolates to series 3, 4 and 5, for assessments of the aggressiveness of pathotype 106 314 0. Isolates were collected in 2012-2013, from Aubusson, Premio or Apache.

| Genotype<sup>a</sup> | Number of isolates | Cultivar tested | Series |
| --- | --- | --- | --- |
| 106 314 0-G1 | 5 | Aubusson | 3+4 |
|  |  | Premio | 3+4 |
|  |  | Michigan Amber | 3+4 |
|  | 1 | Sankara | 5 |
|  |  | Expert | 5 |
|  |  | Bermude | 5 |
| 106 314 0-G2 | 11 | Aubusson | 3+4 |
|  |  | Premio | 3+4 |
|  |  | Michigan Amber | 3+4 |
|  | 2 | Sankara | 5 |
|  |  | Expert | 5 |
|  |  | Bermude | 5 |

<sup>a</sup> Genotyping with 19 SSRs specific to leaf rust (see Table 3)

#### Assessment of aggressiveness

All the phenotyping tests were performed in a greenhouse on eight-day-old wheat seedlings grown in a 21 x 15 x 6 cm plastic box containing potting soil placed under standardized conditions (18°C at night and 22°C during the day, with a 16-h photoperiod). The second leaf of each seedling was maintained, with double-sided tape, on a rigid plate coated with aluminum foil and a 3 cm-long segment of this leaf was inoculated with the fungus. Inoculation was performed with 10 fresh (two-week-old) spores, picked one by one with a human eyelash under a dissecting microscope and deposited on the leaf (Figure 2A and 2B). The plants were then placed in a dew chamber, in the dark, at 15°C, for 18 to 24 h. They were then returned to the greenhouse and the rigid plate was removed. When the first uredinia began to break through the leaf epidermis, generally six days after inoculation, they were counted at 10- to 14-hour intervals (twice daily) until no new uredinia appeared (Figure 2C). Infection efficiency (IE) was estimated for each leaf as the ratio between the final number of uredinia and the number of spores deposited (10 spores). Latency period (LP), estimated as the time between inoculation and the appearance of 50% of the total number of uredinia, was expressed in degree-days, based on the air temperature measured in the greenhouse every ten minutes. Sporulation capacity (SP) was estimated, once the number of uredinia had stabilized, as the number of spores produced per uredinium over a four-day period. Once the final number of uredinia had been reached (i.e. 9 days after inoculation) the spores that had already been produced were removed from the leaf with a small brush. Slightly incurved aluminum gutters (2 x 7cm) made from blind slats were then positioned under each inoculated leaf, which was attached to the gutter with clips (Figure 2D). After four days, all the newly produced spores (Figure 2E) were removed by suction with a cyclone collector into a portion of plastic straw sealed at one end (Figure 2F). Each portion of straw was weighed before and after spore harvesting. SP was calculated by dividing the total weight of the spores collected from a single leaf by the number of uredinia on that leaf.

**Figure 2.**
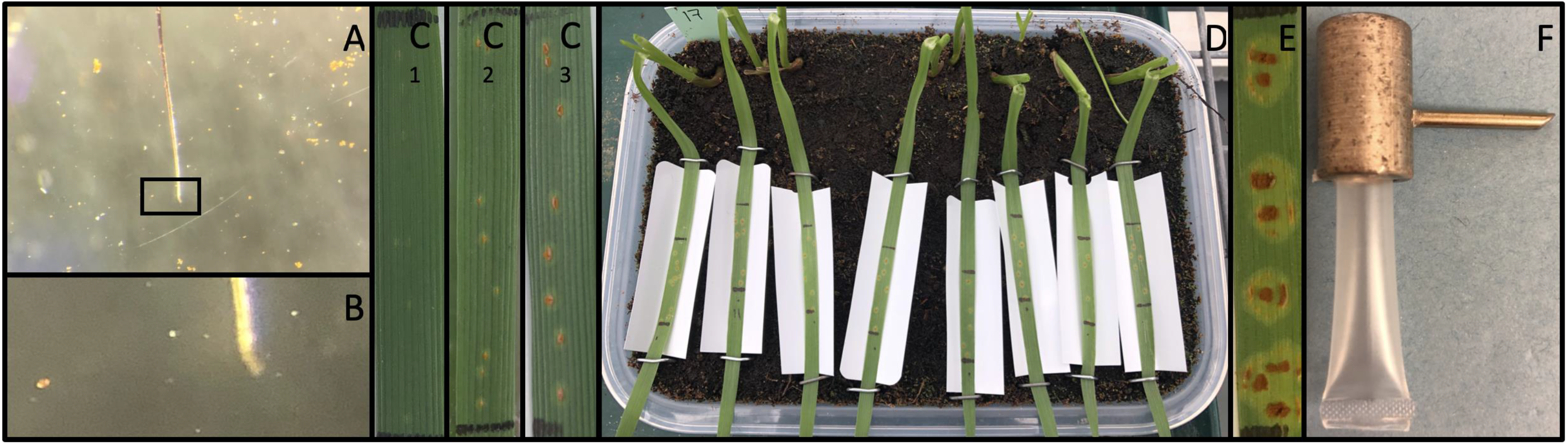
Experimental steps for the assessment of aggressiveness components of *Puccinia triticina* on wheat seedlings. Inoculation: (A) and (B), collection of a spore with a human eyelash and its deposition on a leaf. Latency period (C): (C1) onset of chlorosis, (C2) counting of the uredinia that have broken through the leaf, (C3) end of the latency period, when all the uredinia have emerged. Sporulation: (D) incurved aluminum gutters positioned under the leaves for spore collection, (E) end of sporulation, (F) spores retrieved with a cyclone collector into a sealed portion of plastic straw.

### Statistical analyses

For each of the three aggressiveness traits (the response variable Y), two ANOVA models were used: model (1) for series 1 and 2; model (2) for pooled data from series 3 and 4 (without a series factor, as a preliminary analysis showed that the interactions of the series factor with other factors were never significant), and for series 5. For series 3, 4 and 5, the analysis was performed independently for each cultivar, as the cultivars tested were not the same.

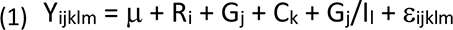

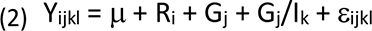

where Y_ijklm_ and Y_ijkl_ are the values of the studied trait in replicate (R) *i*, of genotype (G) *j* and isolate (I) *I* or *k* (isolate being nested within genotype), on cultivar (C) *k* (when this occurred). μ is the overall mean value for this trait and ε is the residual, representing the measurement error, with ε ∼ N(o, σ2)

Log, √ or 1/x transformation was applied to IE, LP and SP when necessary, to obtain a normalized distribution of residuals. When the distribution of residuals could not be normalized by any transformation, a non-parametric Kruskal-Wallis test was performed to analyze the effect of genotype on the aggressiveness components, and also the cultivar effect in series 1 and 2 only. Nevertheless, the Kruskal-Wallis analysis always gave the same results as the ANOVA analysis, even if the data could not be normalized.

All the analyses were performed with R software version 4.1.0.

## RESULTS

### Frequency dynamics of two major *P. triticina* pathotypes in the French landscape in the 2006-2020 period

The frequency of pathotype 106 314 0 in the French *P. triticina* population increased from 30% in 2006 to 48% in 2008 and 51% in 2009 (data from Fontyn *et al*., 2022), the maximum frequency in the landscape for this pathotype (Figure 3). After a plateau at 30-33% from 2011 to 2014, the frequency of this pathotype decreased strongly, to 18% in 2015, 5% in 2016 and less than 1% in 2018. Within pathotype 106 314 0, two main genotypes, 106 314 0-G1 and 106 314 0-G2, were identified and were considered to be dominant as their cumulative frequency ranged from 40% to 65% during the 2006-2016 period (Figure 4). These two genotypes differed at five of the 19 SSR loci studied (RB8, RB11, PtSSR68, PtSSR92 and PtSSR164; Table 3). Genotype 106 314 0-G1 was the most frequent from 2006 to 2012, but it decreased in frequency thereafter (Figure 4). Genotype 106 314 0-G2 was present at a very low frequency from 2007 to 2011. In 2012, the frequencies of the two genotypes were fairly similar, at 23% for 106 314 0-G1 and 30% for 106 314 0-G2. Genotype 106 314 0-G1 then continued to decrease in frequency, eventually disappearing completely from the sampled population in 2014. Conversely, genotype 106 314 0-G2 continued to increase in frequency. It accounted for 42% of the total 106 314 0 pathotype population in 2013 and 66% in 2016. Over the entire period, genotypes other than 106 314 0-G1 and 106 314 0-G2 were also identified (Figure 4; detailed data not shown), which could correspond to other genotypes within pathotype 106 314 0, or to other pathotypes mixed with pathotype 106 314 0 in the original urediniospore bulks.

**Figure 3.**
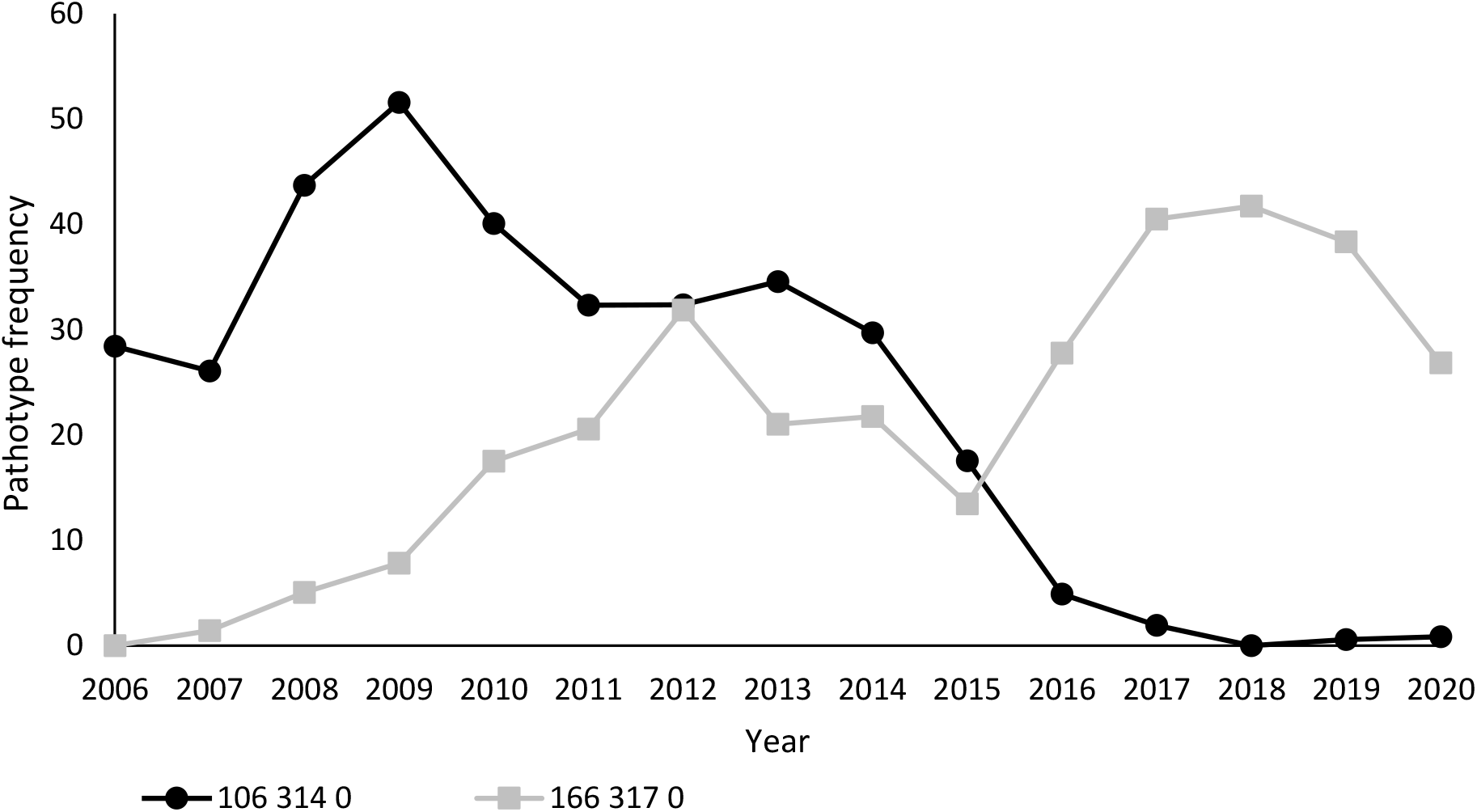
Frequency dynamics of two of the major *Puccinia triticina* pathotypes, 106 314 0 and 166 317 0, in the French landscape during the 2006-2020 period. Pathotype frequency was determined with data from the national survey, which returned 1025 isolates identified as 106 314 0 and 538 isolates identified as 166 317 0 among a total of 3446 pathotyped isolates (data available in Fontyn *et al*., 2022).

**Figure 4.**
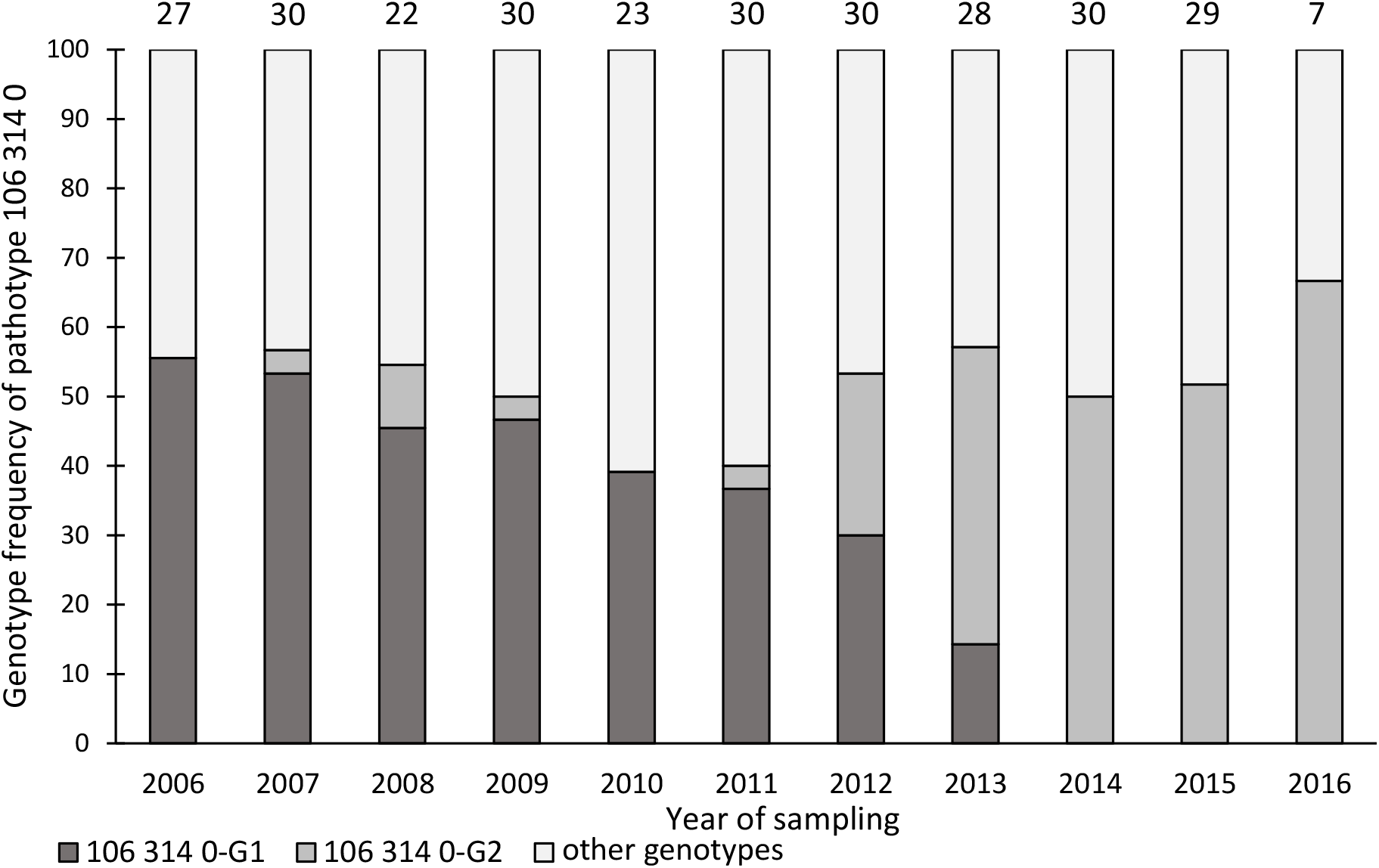
Changes in genotype frequencies within pathotype 106 314 0 in the French landscape during the 2006-2016 period, as determined with 19 SSR markers. The numbers on top of the bars are the numbers of isolates genotyped (286 in total).

**Table 3.** Genotypic characteristics of the two main genotypes, G1 and G2, identified for isolates of pathotypes 166 317 0 and 106 314 0, collected from 2005-2016 for 19 SSR loci. Allele sizes differing between genotypes of the same pathotype are shown in bold typeface.

| Locus | 166 317 0<sup>a</sup> |  |  |  | 106 314 0<sup>a</sup> |  |  |  |
| --- | --- | --- | --- | --- | --- | --- | --- | --- |
|  | 166 317 0-G1 |  | 166 317 0-G2 |  | 106 314 0-G1 |  | 106 314 0-G2 |  |
|  | Allele 1 | Allele 2 | Allele 1 | Allele 2 | Allele 1 | Allele 2 | Allele 1 | Allele 2 |
| RB8 | **143** | **146** | **146** | **152** | **143** | **143** | **143** | **152** |
| RB11 | 200 | 200 | 200 | 200 | **200** | **204** | **175** | **204** |
| RB12 | 282 | 290 | 282 | 290 | 282 | 290 | 282 | 290 |
| RB17 | 190 | 190 | 190 | 190 | 190 | 190 | 190 | 190 |
| RB25 | 230 | 230 | 230 | 230 | 230 | 230 | 230 | 230 |
| RB26 | 352 | 352 | 352 | 352 | 352 | 352 | 352 | 352 |
| PtSSR13 | 129 | 131 | 129 | 131 | 129 | 131 | 129 | 131 |
| PtSSR50 | 365 | 371 | 365 | 371 | 365 | 371 | 365 | 371 |
| PtSSR55 | 308 | 308 | 308 | 308 | 308 | 308 | 308 | 308 |
| PtSSR61 | 296 | 302 | 296 | 302 | 296 | 300 | 296 | 300 |
| PtSSR68 | 309 | 311 | 309 | 311 | **309** | **317** | **309** | **323** |
| PtSSR91 | 380 | 382 | 380 | 382 | 382 | 382 | 382 | 382 |
| PtSSR92 | 246 | 246 | 246 | 246 | **246** | **246** | **246** | **248** |
| PtSSR152 | 389 | 393 | 389 | 393 | 389 | 393 | 389 | 393 |
| PtSSR154 | 247 | 267 | 247 | 267 | 247 | 267 | 247 | 267 |
| PtSSR158 | 235 | 238 | 235 | 238 | 232 | 238 | 232 | 238 |
| PtSSR164 | 219 | 219 | 219 | 219 | **219** | **225** | **219** | **219** |
| PtSSR173 | 213 | 221 | 213 | 221 | 213 | 221 | 213 | 221 |
| PtSSR186 | 341 | 341 | 341 | 341 | 341 | 341 | 341 | 341 |

<sup>a</sup> Seven-digit triplet code (Gilmour, 1973) based on a 20-*Lr* gene differential set: Thatcher lines [*Lr1*, *Lr2a*, *Lr2b*], [*Lr2c*, *Lr3*, *Lr3bg*], [*Lr3ka*, *Lr10*, *Lr13*], [*Lr14a*, *Lr15*, *Lr16*], [*Lr17*, *Lr20*, *Lr23*], [*Lr26*, *Lr17b* (the Australian cv. ‘Harrier’), *Lr37*] and [*Lr24*, *Lr28* (line CS2A/2M)].

Pathotype 166 317 0 first appeared in 2007, initially at a very low frequency (less than 2%). Its temporal evolution in the French *P. triticina* population was characterized by two peaks (data from Fontyn *et al*., 2022). The first peak occurred after a gradual increase of its frequency between 2007 and 2012, reaching 32% of the *P. triticina* population in 2012. Then, it decreased until 2015 to a frequency of 13%, before increasing again to reach a second peak in 2018 at 41% of frequency in the landscape (Figure 3). The frequency of this pathotype finally decreased after 2018 reaching 27% in 2020. Within pathotype 166 317 0, two genotypes, differing by only one of the 19 SSR loci used (RB8; Table 3), were identified and considered to be predominant during the 2013-2016 period. In 2013 and 2014, genotype 166 317 0-G1 dominated, as it accounted for more than 65% of the total 166 317 0 pathotype population (Figure 5). In 2015, genotype 166 317 0-G2 emerged and became co-dominant with 166 317 0-G1. In 2016, 166 317 0-G2 sharply increased in frequency, to 62%, whereas the frequency of 166 317 0-G1 decreased to 7%. Over the four-year period considered, several other genotypes were identified (Figure 5; detailed data not shown), which could correspond to other genotypes within pathotype 166 317 0 or to other pathotypes mixed in the original urediniospore bulks.

**Figure 5.**
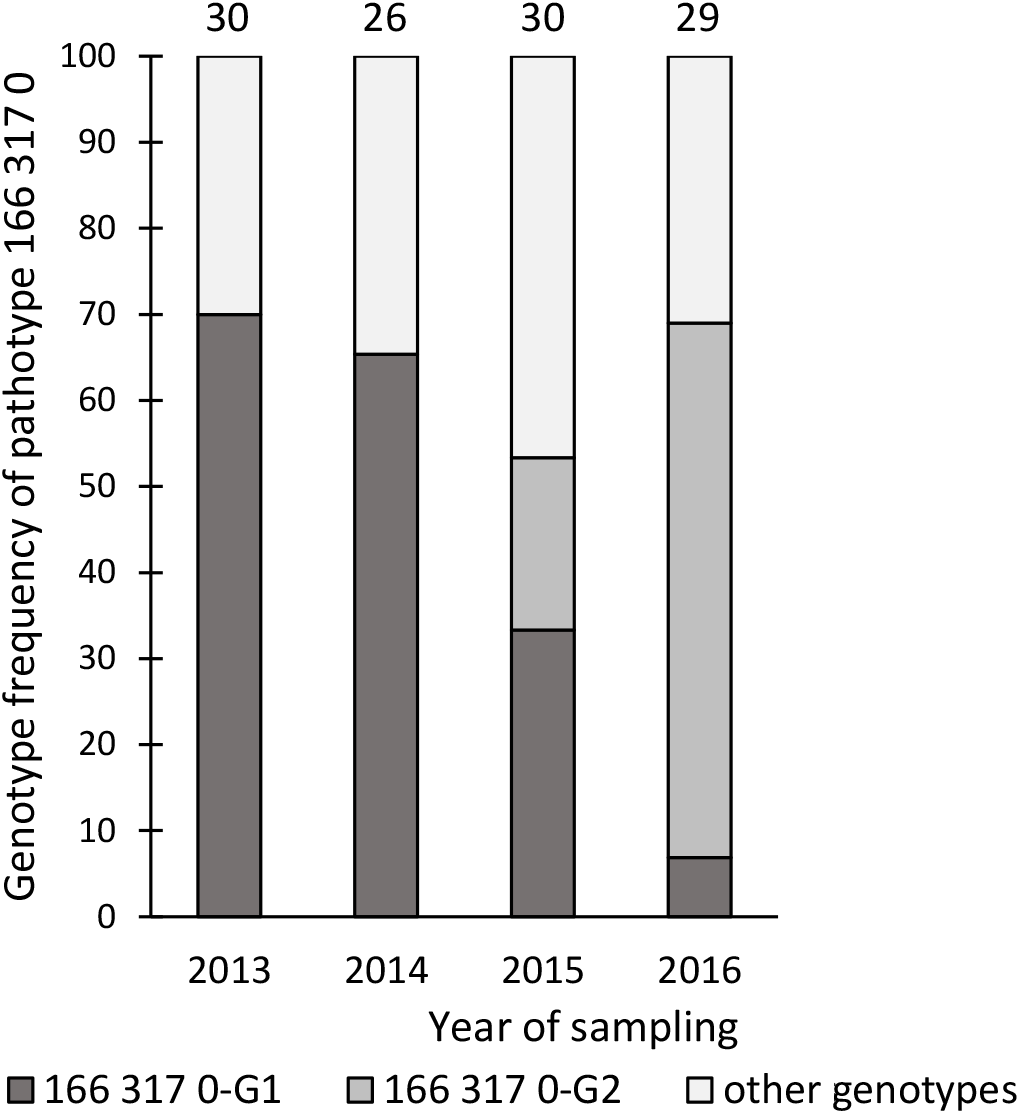
Changes in genotype frequencies within pathotype 166 317 0 in the French landscape during the 2013-2016 period, as determined with 19 SSR markers. Numbers on top of the bars represent the numbers of isolates genotyped (115 in total).

### Differences in aggressiveness between genotypes of the two major *P. triticina* pathotypes expressed on ‘neutral’ and ‘naive’ cultivars

Genotype 166 317 0-G2 was more aggressive than genotype 166 317 0-G1 in assessments of all three aggressiveness components on a ‘neutral’ cultivar, Apache, and a ‘naive’ cultivar Michigan Amber (in analyses of the pooled dataset for the two cultivars; Figure 6A, B and C; Table S5). A comparison of aggressiveness components between the two cultivars is provided in the supplementary data (Figure S1). Infection efficiency (IE) was higher for genotype 166 317 0-G2 (53.1 %) than for genotype 166 317 0-G1 (47.6%). Sporulation capacity (SP) was also higher for genotype 166 317 0-G2, with 0.120 mg/lesion versus 0.114 mg/lesion for genotype 166 317 0-G1. Latency period (LP) was significantly shorter for genotype 166 317 0-G2 (132.6 vs 135.0 degree-days). Significant differences were found between isolates of the two genotypes.

**Figure 6.**
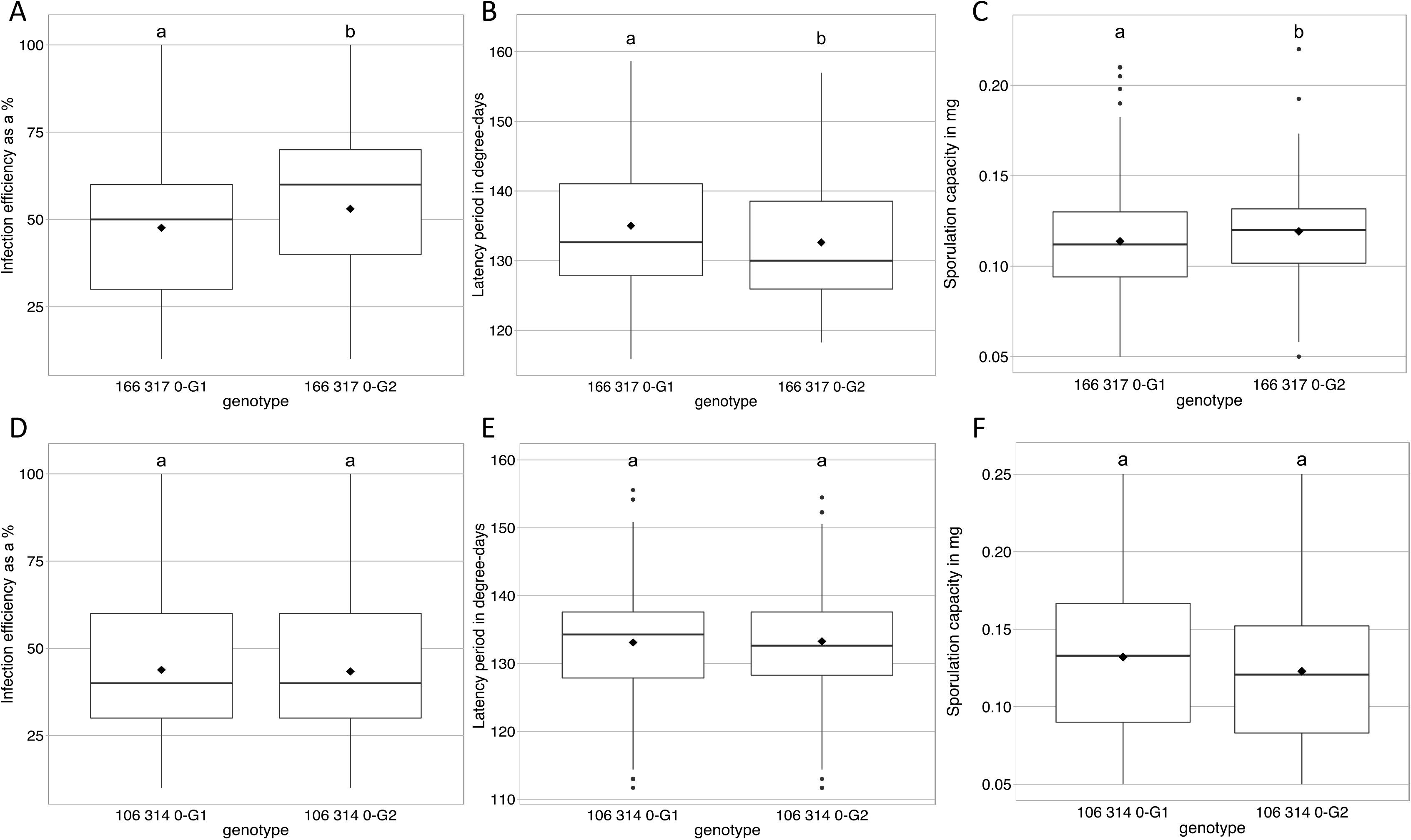
Comparison of genotype aggressiveness within the two major *Puccinia triticina* pathotypes. Infection efficiency (IE) as a %, latency period (LP) in degree-days and sporulation capacity (SP) in mg for pathotypes 166 317 0 (A, B, C) and 106 314 0 (D, E, F) were assessed on cultivars Apache and Michigan (pooled data, see Figures S1 and S2 for results by cultivar). Isolates were sampled from Apache in 2012, 2014, and 2016 for pathotype 166 317 0, and in 2005-2006, 2012-2013 and 2015-2016 for pathotype 106 314 0. Within a box plot, the black diamond represents the mean value and the bar indicates the median value. Letters indicate significant differences between genotypes in Kruskal-Wallis tests (A, C, E, F) or ANOVA (B and D).

IE, LP and SP did not differ between genotypes 106 314 0-G1 and 106 314 0-G2 assessed on cultivars Apache and Michigan Amber (in analyses of the pooled dataset for the two cultivars; Figure 6D, E and F; Table S5). A comparison of aggressiveness components between the two cultivars is provided in the supplementary data (Figure S2). IE for genotype 106 314 0-G1 was 43.8%, which is not significantly different from the value of 43.4% obtained for genotype 106 314 0-G2. SP was 0.128 mg/lesion for genotype 106 314 0-G1, which is not significantly different from the value of 0.124 mg/lesion obtained for genotype G2. Finally, LP differed by less than one degree-day between the two genotypes, at 133.1 degree-days for genotype 106 314 0-G1 and 133.3 degree-days for genotype 106 314 0-G2. By contrast to our findings for pathotype 166 317 0, we observed no significant difference between genotypes of the same pathotype for 106 314 0. This singularity justified our decision to perform phenotypic characterization by comparing the aggressiveness of isolates of pathotype 106 314 0 on other cultivar sets with a view to detecting potential differences.

### Difference in aggressiveness between the two major genotypes of pathotype 106 314 0 on Michigan Amber and major cultivars present in the French varietal landscape

The two major genotypes of pathotype 106 314 0 had significantly different LP on five of the six cultivars tested (Table 4; Table S6). These differences were significant on the cultivar Michigan Amber, whereas no difference was found in analyses of the data for series 1, as described above. IE of genotype 106 314 0-G2 was higher than that of 106 314 0-G1 (48.7% versus 39.1%) on cultivar Expert, but no differences were found on the other cultivars. Genotype 106 314 0-G2 had a shorter LP than 106 314 0-G1 on five of the six cultivars tested. This difference in LP ranged from 2.1 degree-days on Premio to 4.8 degree-days on Expert. A difference of 5 degree-days is equivalent to 8 h at 15°C. Bermude was the only cultivar on which LP did not differ significantly between the two pathogen genotypes. No significant differences in SP were observed between the two genotypes on any of the cultivars tested. SP varied between cultivars, ranging from 0.099 mg on Michigan Amber to 0.134 mg on Sankara, with slightly higher values for 106 314 0-G2 than for 106 314 0-G1. It was not possible to look for significant differences between cultivars, as different cultivars were analyzed in different series.

**Table 4.** Comparison of the aggressiveness of the two major genotypes of pathotype 106 314 0 on five of the most widely grown cultivars in the French landscape during the 2005-2016 period. Letters indicate a significant difference between genotypes on the same cultivar after a Tukey multiple-comparison test or a Wilcoxon-Mann-Whitney test. Numbers in bold typeface indicate a significant difference between isolates within genotypes.

| Cultivar | Genotype | Aggressiveness components |  |  |
| --- | --- | --- | --- | --- |
|  |  | Infection efficiency<sup>z</sup> | Latency period<sup>z</sup> | Sporulation capacity<sup>z</sup> |
| Aubusson | 106 314 0-G1 | 48.5* | **141.0 a** | 0.105 |
|  | 106 314 0-G2 | 49.8* | **137.3 b** | 0.106 |
| Premio | 106 314 0-G1 | 52.6* | **134.5 a** | 0.102 |
|  | 106 314 0-G2 | 49.8* | **132.4 b** | 0.105 |
| Bermude | 106 314 0-G1 | 43.9* | 143.4 | 0.114* |
|  | 106 314 0-G2 | 47.0* | 142.1 | 0.121* |
| Expert | 106 314 0-G1 | 39.1 a | **143.0 a** | 0.120* |
|  | 106 314 0-G2 | 48.7 b | **138.2 b** | 0.132* |
| Sankara | 106 314 0-G1 | 45.0* | 143.9 a* | 0.134* |
|  | 106 314 0-G2 | 46.9* | 140.7 b* | 0.134* |
| Michigan Amber | 106 314 0-G1 | 51.6* | 131.7 a | 0.099 |
|  | 106 314 0-G2 | 52.2* | 129.3 b | 0.102 |

<sup>z</sup> Infection efficiency (IE) was measured as a %, latency period (LP) in degree-days, and sporulation capacity (SP) in mg of spores/uredinia.
\* indicates a significant interaction between genotype and replication factors.

Within genotypes, there was a significant isolate effect for LP, but no significant isolate effect for IE or SP, on Aubusson, Premio and Expert.

For LP, IE and SP, the replication factor (x3) was almost always significant. The interaction between replication factor and the three aggressiveness components was significant for some aggressiveness component measurements (Table 4; Table S6).

## DISCUSSION

Focusing on two major pathotypes from the French *P. triticina* population, we found that several genotypes were present within each pathotype, and that the frequency of the most common genotypes changed over time. The initially dominant genotypes representative of pathotypes 166 317 0 and 106 314 0 were replaced, in each pathotype, by another genotype over the period 2006-2016. The most recent dominant genotype was more aggressive than the older one in both pathotypes.

### Methodological aspects of aggressiveness measurement at the wheat seedling stage

The mean values used to characterize the aggressiveness of different *P. triticina* isolates on wheat seedlings revealed significant variation for each of the three aggressiveness components measured — infection efficiency, latency period and sporulation capacity — at the seedling stage. Aggressiveness and its variation can be measured on seedlings (Milus *et al*., 2006; de Vallavieille-Pope *et al*., 2018) or on adult plants (Lehman & Shaner, 1997; Pariaud *et al*., 2009b; Azzimonti *et al*., 2013). We decided to assess aggressiveness components on wheat seedlings in this study, because the use of seedlings in a semi-controlled environment results in more homogeneous physiological properties of the plant at inoculation and because this approach requires less space and time than studies on adult plants. However, we should be aware that plant stage is known to affect the estimation of the aggressiveness components of *P. triticina* (Pariaud *et al*., 2009b).

With the phenotyping method developed and used in this study, we were able to estimate the three aggressiveness components simultaneously on the same inoculated plant. We measured infection efficiency (IE) more precisely here than in previous studies, in which the variability was sometimes very high. For instance, Pariaud *et al*. (2009b) applied a 1:10 mixture of *P. triticina* urediniospores and *Lycopodium* spores to the leaf surface with a soft brush; they reported IE values ranging from 18% to 80% for the same genotype. This high level of variation resulted from difficulty controlling the number of urediniospores deposited with this method, resulting in differences between experimental series. Although fastidious, the deposition of urediniospores one-by-one on the leaf surface resulted in a much more accurate estimation of IE than other methods based on the dilution of *P. triticina* urediniospores among *Lycopodium* spores or in liquids, such as mineral oils (Pariaud *et al*., 2009b; Sørensen *et al*., 2016). Nevertheless, the interaction between IE and the replication factor was almost always significant, indicating an impact of environmental conditions on this component of aggressiveness. This interaction reflects the difficulty ensuring uniform dew quality in the dew chamber just after inoculation, as the success of urediniospore germination and penetration depends on the presence of a water film on the leaf surface (Bolton *et al*., 2008).

Sporulation capacity (SP) was measured by collecting urediniospores produced between 9 and 12 days after inoculation, and differences in SP between genotypes were detected for only one experimental series. Pariaud *et al*. (2009b) found differences in SP between pathotypes of *P. triticina* in a study in which urediniospores were collected 15 to 23 days after the inoculation of adult plants. They also found that the difference in SP between isolates increased with the regular collection of urediniospores until 59 days after inoculation. The collection of urediniospores more than 13 days after inoculation is of potential relevance for future experiments with our method, to maximize differences and reveal small differences in this component of aggressiveness between isolates.

Latency period (LP) was the component that differed most between pathogen genotypes and was the least affected by the replication factor. The calculation of LP in degree-days made it possible to take the temperature-dependence of both infection processes and pathogen development in the leaf into account (Lovell *et al*., 2004). In most pathosystems, LP is a component of choice for studies of quantitative interactions (aggressiveness, partial resistance) because time is a variable that is much easier to fractionate and, therefore, to quantify precisely, than any biological trait, making it possible to highlight extremely small differences between isolates repeatedly. This is almost an epistemological issue for experimental epidemiology and phytopathometry (Suffert & Thompson, 2018; Bock *et al*., 2022).

### Evolution of greater aggressiveness in two *P. triticina* pathotypes

In the case of pathotype 166 317 0, genotype 166 317 0-G2 was more aggressive than the one it replaced, 166 317 0-G1, with highly consistent results obtained for the different components of aggressiveness. Following this switch between genotypes, the frequency of pathotype 166 317 0 increased considerably over a period of three years, whereas the proportion of compatible host cultivars in the landscape decreased (data not shown). Severe *P. triticina* epidemics have been shown to be associated with a high SP, a high IE and a short LP (Azzimonti *et al*., 2022). In pathotype 106 314 0, comparisons of aggressiveness on a ‘neutral’ cultivar revealed no significant difference between the most recent and oldest genotypes. A comparison of the aggressiveness of these two genotypes on some of the cultivars most frequently grown in the landscape revealed that 106 314 0-G2 was more aggressive than 106 314 0-G1 only for LP. This change in predominant genotype coincided with a halt in the decline of pathotype 106 314 0 frequency in the landscape. The shorter LP of genotype 106 314 0-G2 seems to have an effect on the frequency of this pathotype in the landscape, consistent with the assertions of several studies that LP is the aggressiveness component with the largest effect on pathogen dynamics in field conditions (Lannou, 2012). In modeling studies, this also appeared to be the trait with the largest impact on the intensity of *P. triticina* epidemics, as it determined the number of reproductive cycles (i.e. from inoculation to spore dispersion) of the pathogen possible in a single season (Rimbaud *et al*., 2018). Although only LP differed significantly between 106 314 0-G2 and 106 314 0-G1, the data for the other aggressiveness components also supported the notion that 106 314 0-G2 was more aggressive, for every component, on all cultivars, except for IE on Premio. Measurements of aggressiveness components on seedlings in controlled conditions (only one reproductive cycle of the pathogen) have a limited capacity for the detection of small phenotypic variations.

The competitive advantage of one particular genotype within a pathotype, expressing a difference in parasitic fitness for the same virulence profile, may account for the replacement of one genotype with another. This hypothesis is particularly realistic here as the emergence of the more aggressive genotype in each pathotype, 106 314 0 and 166 317 0, coincided with a short-term change in pathotype frequency: (i) the decrease in the frequency of pathotype 106 314 0 in the landscape observed from 2009 to 2011 was temporarily halted during the 2011-2014 period, when the frequency of this pathotype reached a plateau (Figure 3), coinciding with the replacement of the less aggressive genotype 106 314 0-G1 with the more aggressive genotype 106 314 0-G2 in 2012 and 2013 (Figure 4); (ii) the decrease in the frequency of pathotype 166 317 0 in the landscape observed from 2012 to 2015 was halted over the 2015-2018 period, resulting in a new peak frequency (Figure 3) coinciding with the replacement of the less aggressive 166 317 0-G1 genotype with the more aggressive 166 317 0-G2 genotype in 2012 and 2013 (Figure 5).

These results suggest that the replacement of some pathotypes by others, driven by changes in the frequencies of resistance genes in the varietal landscape, may be slowed, to various extents, by increases in the aggressiveness of certain genotypes or ‘lineages’ (defined as genetically related genotypes of the same group of pathotypes, i.e. with the same close ancestor), consistent with the recent analysis by Fontyn *et al*. (2022). It is even possible, as for pathotype 166 317 0, for the pathotype to gain a ’second life’ — for a period of three to four years — due to an increase in the aggressiveness of a new genotype, as long as it remains adapted (or at least not maladapted) to the resistance genes present in the varieties deployed in the landscape. This empirical study shows that differences in aggressiveness can be expressed on a ‘neutral’ cultivar (for pathotype 166 317 0), but sometimes (for pathotype 106 314 0) only on ‘non-neutral’ cultivars representative of a varietal landscape. This finding clearly complicates the experimental approach (difficulty determining the most appropriate experimental design without making assumptions), but also the data analysis. The magnitude of the differences in aggressiveness probably depends on the type of cultivars tested (‘neutral’ or ‘non-neutral’). Indeed, we found that the intensity of the mid-term changes in frequencies highlighted above was variable, with either a simple slowing of a trend towards a decrease (plateau, as for pathotype 106 314 0), or a change in direction, with a new increase (peak, as for pathotype 166 317 0).

### Putative origin of the more aggressive genotypes

All surveys in wheat-growing areas to date have provided evidence for continual evolution of *P. triticina* populations, with rapid changes in pathotype frequencies (Goyeau *et al*., 2012; Kosman *et al*., 2019; Zhang *et al*., 2020; Fontyn *et al*., 2022; Kolmer & Fajolu, 2022). These changes can be explained intrinsically by the acquisition of new virulences, by mutations, somatic exchanges or, more rarely, genetic recombination. A study on the Australian *P. striiformis* f. sp. *tritici* population showed that it had evolved due to mutation events (Steele *et al*., 2001). Whole-genome sequencing has detected similar recurrent mutations in *P. triticina*, suggesting that such events play a major role in genetic variability within clonal lineages (Fellers *et al*., 2021). Somatic exchange may also have played a role, as in the emergence of the Ug99 lineage of *Puccinia graminis* f. sp. *tritici,* which had one identical haploid nucleus in common with an old African lineage, with no sign of recombination (Li *et al*., 2019). A similar situation has been described in *P. triticina*, with the Australian isolate Pt64 resulting from somatic exchange between two parental isolates (Wu *et al*., 2019). More rarely, genetic recombination may also underlie the emergence of new genotypes with new virulences. Indeed, *P. triticina* is a heteroecious fungus that requires an alternative host, *Thalictrum speciosissimum* (not naturally present in most places worldwide, including France), for sexual reproduction. The high proportion of repeated genotypes and the heterozygosity rates of European and French leaf rust populations confirmed the very dominant role of clonal reproduction (Goyeau *et al*., 2007; Kolmer *et al*., 2013). Genetic changes do not necessarily result in different virulence profiles, so isolates with identical or very similar pathotypes may have different origins. In a previous study on *P. triticina* nine microsatellite markers were used to analyze the genotypes of 33 pathotypes (Goyeau *et al*., 2007). This analysis highlighted the presence of different genotypes, differing by only one of the nine SSRs tested, in five of the pathotypes. Another study on 121 European *P. triticina* isolates with 21 SSR markers revealed a significant correlation between phenotype and genotype, but the same phenotype could, nevertheless, be associated with several genotypes (Kolmer *et al*., 2013). The results of our study raise questions as to the genetic relationship between the most recent genotypes, 166 317 0-G2 and 106 314 0-G2, and the oldest genotypes, 166 317 0-G1 and 106 314 0-G1, respectively. Can the most recent and the oldest genotypes be considered to belong to the same ‘lineage’, as previously defined? A mutation event during asexual reproduction probably led to the emergence of the most recent genotype (G2) of pathotype 166 317 0, which differs from the older genotype (G1) by only one SSR allele (Table 3). By contrast, genotypes G1 and G2 of pathotype 106 314 0 are more differentiated, as they are differing by five SSR alleles (Table 3). Genotype 106 314 0-G2 may, therefore, be the result of several mutation events occurring in the same lineage and conferring greater aggressiveness than for 106 314 0-G1. Alternatively, the presence of 106 314 0-G2 in France may reflect the introduction from an exogenous source of a different (more aggressive) lineage that had acquired the same virulences and, therefore, belonged to the same pathotype, 106 314 0. Migration events are frequent in cereal rust populations, as already shown for yellow rust (Ali *et al*., 2014; Bueno-Sancho *et al*., 2017), for which there is a famous example of migration in the form of an introduction into Australia from northwest Europe in 1979 (Wellings & McIntosh, 1990). Studies on *P. triticina* in wheat-growing areas worldwide have revealed a broad geographic distribution of identical and highly related multilocus genotypes, highlighting the potential of leaf rust for long-distance migration (Kolmer *et al*., 2019) and the possibility of new genotypes resulting from exogenous introduction.

### Additional approaches for detecting genetic variation related to aggressiveness

This study was not designed to characterize overall pathotypic or genotypic changes in the *P. triticina* population, so our data should be extrapolated with caution, as any attempt to use them for such purposes would be subject to multiple sampling biases. In addition to the four genotypes on which we focused here, several other genotypes (with cumulative frequencies of 31% to 61%, depending on the year) were identified. This is due to the initial pathotyping performed annually for the national survey and the genotyping for this study being performed on different isolates, purified from a bulk of urediniospores collected from a single leaf, which may well have been infected with several pathotypes and/or genotypes (Figure 1). This highlights the imperfections of the protocol linked to the constraints of working on datasets and biological material acquired over several decades with ever-changing techniques. It would have been better to pathotype and phenotype the 401 isolates from the same purified batch of urediniospores, as we did later on for the 44 isolates for which aggressiveness was characterized (Figure 1). However, given the size of the sample, this limitation did not prevent us from obtaining a correct overview of changes for the most common genotypes.

Our results revealed rapid genotype evolution within the pathotypes that was not detectable if only virulence phenotypes were considered. These findings demonstrate the value of analyzing population dynamics not only from the pathotype standpoint, but also with genotype data, to obtain a more informative picture of pathogen diversity. For leaf rust, as for many fungal pathogens that are commonly described by their virulence profile, there is no univocal link between the pathotypic and the genotypic characterization. In practice, pathotypic characterization and naming has been favoured because of the importance of virulence in the structure of populations. Our results show that different genotypes (defined here by differences in the combination of SSR markers) that have the same virulence profile can nevertheless express differences in the quantitative component of pathogenicity (aggressiveness). Such differences are potentially related to interactions with sources of quantitative resistance. Conversely, identical genotypes may differ in one or several virulences. In summary, differentiation (or not) for neutral markers is independent from differentiation (or not) for functional mutations. There is currently no nomenclature for the designation of a unique association between one pathotype and one genotype. Therefore, we propose to define a unique association with the term ‘pathogenotype’, exemplified here with 106 314 0-G1 and 106 314 0-G2.

The use of a small number of SSR markers is not sufficient for the detection of genetic variations in *P. triticina* populations. Genome-wide genotyping approaches are required to characterize the genetic diversity of leaf rust populations more precisely and to explain the emergence of new genotypes. As an illustration of this approach, a combined genome and transcriptome analysis has been performed on 133 *P. striiformis* f. sp. *tritici* isolates collected in 16 European countries. This analysis provided more precise information about the origin of the new emerging races, and showed that SNP analysis was an effective approach for the detection of pathogen diversity and for pathogen surveillance (Bueno-Sancho *et al*., 2017). Similarly, Fellers *et al*. (2021) genotyped 121 *P. triticina* isolates with 121 907 SNP markers, and showed that recurrent mutation and selection had played a major role in differentiation within clonal lineages. The results of our study highlight the importance of combining genome-wide genotyping tools with precise pathotyping and aggressiveness phenotyping to detect the emergence of new variants and to improve our understanding of population dynamics.

## Acknowledgments

We thank Nathalie Retout for technical assistance in the preparation of aggressiveness experiments. We thank the French Wheat Breeders groups ‘Recherches Génétiques Céréales’ and ‘CETAC’, and ‘ARVALIS-Institut du Végétal’ for their help collecting strains of *P. triticina* from their nurseries and field trials. We thank Dr. Julie Sappa for editing the manuscript.

## Data Availability Statement

The data that support the findings of this study are openly available in the INRAE Dataverse online data repository (https://data.inrae.fr/) at https://doi.org/10.57745/MZ8TDK

