## Supplementary Material for "Evolution within a given virulence phenotype (pathotype) is driven by changes in aggressiveness: a case study of French wheat leaf rust populations" (from another version of this preprint: version 3, 2023-03-01)

### SUPPLEMENTARY TABLES

**Table S1.** Number of isolates identified as pathotype 106 314 0 collected on 14 of the most frequently grown bread wheat cultivars during the 2006-2016 period and genotyped with 19 SSRs (see Table 3).

| | | | Year of sampling | | | | | | | | | | | |
|---|---|---|---|---|---|---|---|---|---|---|---|---|---|---|
| Cultivar | Registration year | Postulated resistance genes[a] | 2006 | 2007 | 2008 | 2009 | 2010 | 2011 | 2012 | 2013 | 2014 | 2015 | 2016 | Total |
| Charger | 1997 | *Lr10, Lr13* | 4 | | | | | | | | | | | 4 |
| Tremie | 1992 | *Lr10, Lr13* | 5 | | | | | | | | | | | 5 |
| Sankara | 2004 | *Lr10, Lr13, Lr37* | 4 | 5 | 3 | 5 | | | | | | | | 17 |
| Aubusson | 2002 | *Lr10, Lr13, Lr37* | 5 | 5 | 3 | 5 | 3 | 5 | 5 | | | | | 31 |
| Apache | 1998 | *Lr13, Lr37* | 9 | 10 | 9 | 10 | 8 | 10 | 10 | 9 | 10 | | | 85 |
| Orvantis | 2000 | *Lr10, Lr13, Lr37* | | 5 | | | | | | | | | | 5 |
| Soissons | 1988 | *Lr14a* | | 5 | 4 | 5 | | | | | | | | 14 |
| Mendel | 2004 | *Lr13* | | | 3 | | | | | | | | | 3 |
| Premio | 2007 | *Lr14a, Lr37* | | | | 5 | 4 | 5 | | | | | | 14 |
| Bermude | 2007 | *Lr10, Lr13, Lr14a, Lr37* | | | | | 4 | 5 | 5 | 5 | | 7 | | 26 |
| Arezzo | 2008 | *Lr10, Lr14a, Lr37* | | | | | 4 | 5 | 5 | 5 | 5 | 8 | 1 | 33 |
| Expert | 2008 | *Lr1, Lr13* | | | | | | | 5 | 4 | 5 | | 3 | 17 |
| Pakito | 2011 | *Lr13, Lr37* | | | | | | | | 5 | 5 | 7 | | 17 |
| Solehio | 2009 | *Lr14a, Lr37* | | | | | | | | | 5 | 7 | 3 | 15 |
| Number of isolates | | | 27 | 30 | 22 | 30 | 23 | 30 | 30 | 28 | 30 | 29 | 7 | 286 |

[a] *Lr* genes were postulated by performing multipathotype tests with standard isolates with known virulence genes (Fontyn *et al.*, 2022)

**Table S2.** Number of isolates identified as pathotype 166 317 0 collected on eight of the most frequently grown bread wheat cultivars during the 2013-2016 period and genotyped with 19 SSRs (see Table 3).

| | | | Year of Sampling | | | | |
|---|---|---|---|---|---|---|---|
| Cultivar | Registration year | Postulated resistance genes[a] | 2013 | 2014 | 2015 | 2016 | Total |
| Altigo | 2007 | *Lr3, Lr13, Lr37* | 8 | | | | 8 |
| Bermude | 2007 | *Lr10, Lr13, Lr14a, Lr37* | 5 | | 5 | | 10 |
| Arezzo | 2008 | *Lr10, Lr14a, Lr37* | 7 | 9 | 5 | 10 | 31 |
| Pakito | 2011 | *Lr13, Lr37* | 5 | 4 | 5 | | 14 |
| Solehio | 2009 | *Lr14a, Lr37* | 5 | 5 | | 5 | 15 |
| Cellule | 2011 | *Lr3* | | 8 | 10 | 9 | 27 |
| Apache | 1998 | *Lr13, Lr37* | | | 5 | | 5 |
| Fructidor | 2014 | *Lr13, Lr14a* | | | | 5 | 5 |
| Number of isolates | | | 30 | 26 | 30 | 29 | 115 |

[a] *Lr* gnes were postulated by performing multipathotype tests with standard isolates with known virulences (Fontyn *et al.*, 2022)

**Table S3.** Experimental design for assessments of the aggressiveness of the two main genotypes (G1 and G2) of each of the two major pathotypes (106 314 0 and 166 317 0) of *Puccinia triticina* collected in France during 2005-2016: distribution of the isolates over five experimental series, cultivar from which the isolate was collected, and cultivars on which the aggressiveness of the isolate was assessed.

| Isolate | Pathotype | Genotype | Series | Cultivar sampled | Cultivars tested |
|---|---|---|---|---|---|
| BT12M202 | 166 317 0 | 166 317 0-G1 | 2 | Apache | Apache, Michigan Amber |
| BT12M378 | 166 317 0 | 166 317 0-G1 | 2 | Apache | Apache, Michigan Amber |
| BT12M109 | 166 317 0 | 166 317 0-G1 | 2 | Apache | Apache, Michigan Amber |
| BT12M391 | 166 317 0 | 166 317 0-G1 | 2 | Apache | Apache, Michigan Amber |
| BT12M192 | 166 317 0 | 166 317 0-G1 | 2 | Apache | Apache, Michigan Amber |
| BT12M321 | 166 317 0 | 166 317 0-G1 | 2 | Apache | Apache, Michigan Amber |
| BT14M018 | 166 317 0 | 166 317 0-G1 | 2 | Apache | Apache, Michigan Amber |
| BT14M066 | 166 317 0 | 166 317 0-G1 | 2 | Apache | Apache, Michigan Amber |
| BT14M149 | 166 317 0 | 166 317 0-G1 | 2 | Apache | Apache, Michigan Amber |
| BT14M254 | 166 317 0 | 166 317 0-G1 | 2 | Apache | Apache, Michigan Amber |
| BT14M344 | 166 317 0 | 166 317 0-G1 | 2 | Apache | Apache, Michigan Amber |
| BT14M392 | 166 317 0 | 166 317 0-G1 | 2 | Apache | Apache, Michigan Amber |
| BT16M219 | 166 317 0 | 166 317 0-G2 | 2 | Apache | Apache, Michigan Amber |
| BT16M164 | 166 317 0 | 166 317 0-G2 | 2 | Apache | Apache, Michigan Amber |
| BT16M246 | 166 317 0 | 166 317 0-G2 | 2 | Apache | Apache, Michigan Amber |
| BT16M174 | 166 317 0 | 166 317 0-G2 | 2 | Apache | Apache, Michigan Amber |
| BT05M021 | 106 314 0 | 106 314 0-G1 | 1 | Apache | Apache, Michigan Amber |
| BT05M241 | 106 314 0 | 106 314 0-G1 | 1 | Apache | Apache, Michigan Amber |
| BT05M067 | 106 314 0 | 106 314 0-G1 | 1 | Apache | Apache, Michigan Amber |
| BT06M138 | 106 314 0 | 106 314 0-G1 | 1 | Apache | Apache, Michigan Amber |
| BT06M195 | 106 314 0 | 106 314 0-G1 | 1 | Apache | Apache, Michigan Amber |

| | | | | | |
|---|---|---|---|---|---|
| BT06M101 | 106 314 0 | 106 314 0-G1 | 1 | Apache | Apache, Michigan Amber |
| BT15M292 | 106 314 0 | 106 314 0-G2 | 1 | Apache | Apache, Michigan Amber |
| BT15M011 | 106 314 0 | 106 314 0-G2 | 1 | Apache | Apache, Michigan Amber |
| BT16V048 | 106 314 0 | 106 314 0-G2 | 1 | Apache | Apache, Michigan Amber |
| BT15M121 | 106 314 0 | 106 314 0-G2 | 1 | Apache | Apache, Michigan Amber |
| BT15M309 | 106 314 0 | 106 314 0-G2 | 1 | Apache | Apache, Michigan Amber |
| BT15M035 | 106 314 0 | 106 314 0-G2 | 1 | Apache | Apache, Michigan Amber |
| BT12M380 | 106 314 0 | 106 314 0-G1 | 3 | Aubusson | Aubusson, Premio, Michigan Amber |
| BT12M355 | 106 314 0 | 106 314 0-G1 | 3 | Premio | Aubusson, Premio, Michigan Amber |
| BT12M137 | 106 314 0 | 106 314 0-G1 | 3 | Premio | Aubusson, Premio, Michigan Amber |
| BT13V142 | 106 314 0 | 106 314 0-G2 | 3 | Aubusson | Aubusson, Premio, Michigan Amber |
| BT13M108 | 106 314 0 | 106 314 0-G2 | 3 | Aubusson | Aubusson, Premio, Michigan Amber |
| BT13V026 | 106 314 0 | 106 314 0-G2 | 3 | Aubusson | Aubusson, Premio, Michigan Amber |
| BT13M168 | 106 314 0 | 106 314 0-G2 | 3 | Aubusson | Aubusson, Premio, Michigan Amber |
| BT12M281 | 106 314 0 | 106 314 0-G2 | 3 | Aubusson | Aubusson, Premio, Michigan Amber |
| BT12M302 | 106 314 0 | 106 314 0-G2 | 3 | Premio | Aubusson, Premio, Michigan Amber |
| BT12M379 | 106 314 0 | 106 314 0-G2 | 3 | Premio | Aubusson, Premio, Michigan Amber |
| BT12M284 | 106 314 0 | 106 314 0-G2 | 3 | Premio | Aubusson, Premio, Michigan Amber |
| BT12M119 | 106 314 0 | 106 314 0-G1 | 1+4 | Apache | Aubusson, Premio, Michigan Amber, Apache |
| BT12M033 | 106 314 0 | 106 314 0-G2 | 1+4 | Apache | Aubusson, Premio, Michigan Amber, Apache |
| BT13V137 | 106 314 0 | 106 314 0-G1 | 1+4+5 | Apache | Aubusson, Premio, Michigan Amber, Sankara, Expert, Bermude, Apache |
| BT13V189 | 106 314 0 | 106 314 0-G2 | 1+4+5 | Apache | Aubusson, Premio, Michigan Amber, Sankara, Expert, Bermude, Apache |
| BT12M326 | 106 314 0 | 106 314 0-G2 | 1+4+5 | Apache | Aubusson, Premio, Michigan Amber, Sankara, Expert, Bermude, Apache |

**Table S4.** Allocation of 19 SSR markers to two multiplexes, for genotyping *Puccinia triticina* isolates.

| | SSR ID | Fluorescence label | Forward primer | Reverse primer | Chromosome position | Allele size range | Repeat size | Reference |
|---|---|---|---|---|---|---|---|---|
| Multiplex 1 | **RB8** | Vic | CGCCGTTCCCATCGTTC | TAAAACACTCCACCCACGCC | 17A/17B | 143-155 | 3 | Duan et al., 2003 |
| | **PtSSR173** | Vic | CTCAGCGACCTCAAAGAACC | GAGACGACGGATGTCAACAA | 2A/2B | 213-223 | 2 | Szabo & Kolmer, 2007 |
| | **PtSSR186** | Vic | GCCACGAGAAATACATAGAAATAAAA | GGTTGTTGATGGGCTTGAGT | 10A/10B | 338-353 | 3 | Szabo & Kolmer, 2007 |
| | **RB11** | Pet | AGCAGTGAGCAGCAGCGTC | ACTACTGTGAGTGTCGGCTTGG | - | 175-206 | 2 | Duan et al., 2003 |
| | **PtSSR158** | 6Fam | GACGACTTCGTCACTGCTGA | GAGGAGAAGCCGTTCTGTTG | 3A/3B | 226-238 | 3 | Szabo & Kolmer, 2007 |
| | **PtSSR68** | Pet | GACTCAGCCCACTGCTAACC | GATGGCGACGTATTTGGTCT | 9A/9B | 309-341 | 2 | Szabo & Kolmer, 2007 |
| | **PtSSR50** | 6Fam | CATCGGAATGGTCTGTCTCC | CCAAATGCTATGAGTGGAAAA | 12A/12B | 365-373 | 2 | Szabo & Kolmer, 2007 |
| | **PtSSR164** | Ned | GTGGAAGTGAGCGGAAGAAG | GGAGATGGGCAGATGAGGTA | 10A/10B | 219-233 | 2 | Szabo & Kolmer, 2007 |
| | **PtSSR61** | Ned | CGAACTGGTACAACGCACTG | CGCAAAAAGGCTGATCTCTG | 17A/17B | 296-302 | 2 | Szabo & Kolmer, 2007 |
| Multiplex 2 | **RB17** | Vic | CTTCGGTAGGATTTCGAGCG | CAGCTCCAAATCCTTTGCC | 14A/14B | 190-193 | 3 | Duan et al., 2003 |
| | **PtSSR92** | Pet | CCAAGGAACAGTCCACCAAG | GAGTCGGGTAAGCCATCTGA | 1A/1B | 246-258 | 2 | Szabo & Kolmer, 2007 |
| | **RB12** | Vic | CCACAAGCAACCACATACCACC | TGGTCCATGAAGAAGTCTCTGAAC | 16A/16B | 282-290 | 2 | Duan et al., 2003 |
| | **RB26** | Vic | TCGTCCTGCCTACCTCTGAC | AAAGTGCATGATCTGCATGTG | 16A/16B | 349-352 | 3 | Duan et al., 2003 |
| | **PtSSR91** | Ned | ATCTTGCGTCTCAGCCATCT | CGCCGCTCTTCATCTCTTAC | 1A/1B | 380-384 | 2 | Szabo & Kolmer, 2007 |
| | **PtSSR13** | 6Fam | CGAATTCGCGTTTTATGTCC | TGATCCAATCGAACCTAGCC | 2A/2B | 129-131 | 2 | Szabo & Kolmer, 2007 |
| | **RB25** | 6Fam | ATGTCTGTAGTCGGCAGGGC | GCCTCTGCGGGATCGGT | 4A/4B | 228-230 | 2 | Duan et al., 2003 |
| | **PtSSR55** | 6Fam | AGCTTACGGTCCTCAATCG | AGTGAAAGGGGCTGGGAGT | 18A/18B | 308-310 | 2 | Szabo & Kolmer, 2007 |
| | **PtSSR152** | 6Fam | CTCCGTTCCTCTTTCTGTCG | CCATCGCAACCAACAAACA | 18A/18B | 389-393 | 2 | Szabo & Kolmer, 2007 |
| | **PtSSR154** | Ned | ACGGTCAACAGCCAACTACC | CCTCGTCATCCTGGTTGAGT | 16A/16B | 247-276 | 3 | Szabo & Kolmer, 2007 |

**Table S5.** ANOVA *p-values* for series 1 and 2 testing whether the two genotypes (106 314 0-G1 and 106 314 0-G2 ; 166 317 0-G1 and 166 317 0-G2) within each *P. triticina* pathotype (106 314 0 and 166 317 0) differ in the aggressiveness components infection efficiency (IE), latency period (LP), and sporulation capacity (SP), on pooled data from cutivars Apache and Michigan.

| | | | Source of variation | | | | | | | |
|---|---|---|---|---|---|---|---|---|---|---|
| Series | Pathotype | Component[1] | Repetition | df | Genotype | df | Cultivar | df | Isolate/Genotype | df |
| | | IE | <0.0001 | 2 | 0.9 | 1 | 0.1 | 1 | <0.0001 | 15 |
| 1 | 106 314 0 | LP | <0.0001 | 2 | 0.7 | 1 | <0.0001 | 1 | 0.2 | 15 |
| | | SP | <0.0001 | 2 | 0.05 | 1 | 0.03 | 1 | <0.0001 | 15 |
| | | IE | <0.0001 | 2 | 0.0001 | 1 | 0.0002 | 1 | <0.0001 | 14 |
| 2 | 166 317 0 | LP | <0.0001 | 2 | 0.0003 | 1 | <0.0001 | 1 | 0.01 | 14 |
| | | SP | 0.003 | 2 | 0.01 | 1 | 0.004 | 1 | 0.0002 | 14 |

[1] An underlined component means that the data did not follow a normal distribution required for the ANOVA, however a non-parametric test (Kruskal-Wallis) gave the same results.

**Table S6.** ANOVA *p-values* for series 3, 4 and 5 testing whether the two genotypes 106 314 0-G1 and 106 314 0-G2 differ in the aggressiveness components infection efficiency (IE), latency period (LP), and sporulation capacity (SP), as assessed on Michigan and on five of the most widely grown cultivars in France during the 2005-2016 period.

| | | | Source of variation | | | | | |
|---|---|---|---|---|---|---|---|---|
| Series | Tested cultivar | Component[1] | Repetition | df | Genotype[2] | df | Isolate/Genotype | df |
| 3+4 | Aubusson | IE | <0.0001 | 2 | **0.8** | 1 | 0.1 | 14 |
| | | LP | 0.001 | 2 | <0.0001 | 1 | <0.0001 | 14 |
| | | SP | 0.03 | 2 | **0.9** | 1 | <0.0001 | 14 |
| 3+4 | Premio | IE | <0.0001 | 2 | **0.2** | 1 | 0.09 | 14 |
| | | LP | <0.0001 | 2 | 0.0006 | 1 | 0.001 | 14 |
| | | SP | 0.002 | 2 | **0.8** | 1 | <0.0001 | 14 |
| 3+4 | Michigan | IE | <0.0001 | 2 | **1** | 1 | 0.6 | 14 |
| | | LP | 0.008 | 2 | <0.0001 | 1 | 0.2 | 14 |
| | | SP | <0.0001 | 2 | 0.1 | 1 | <0.0001 | 14 |
| 5 | Bermude | IE | 0.0005 | 2 | **0.5** | 1 | 0.2 | 1 |
| | | LP | 0.1 | 2 | 0.4 | 1 | 1 | 1 |
| | | SP | <0.0001 | 2 | **0.4** | 1 | 0.1 | 1 |
| 5 | Expert | IE | 0.0001 | 2 | 0.02 | 1 | 0.6 | 1 |
| | | LP | 0.12 | 2 | 0.003 | 1 | 0.03 | 1 |
| | | SP | 0.01 | 2 | **0.1** | 1 | 0.7 | 1 |
| 5 | Sankara | IE | <0.0001 | 2 | **0.7** | 1 | 0.1 | 1 |
| | | LP | <0.0001 | 2 | **0.002** | 1 | 0.2 | 1 |
| | | SP | <0.0001 | 2 | **0.9** | 1 | 0.3 | 1 |

[1] An underlined component means that the data did not follow a normal distribution required for the ANOVA, however a non-parametric test (Kruskal-Wallis) gave the same results.

[2] *p-values* in bold indicate a significant interaction effect between the repetition factor and the component.

**SUPPLEMENTARY FIGURES**

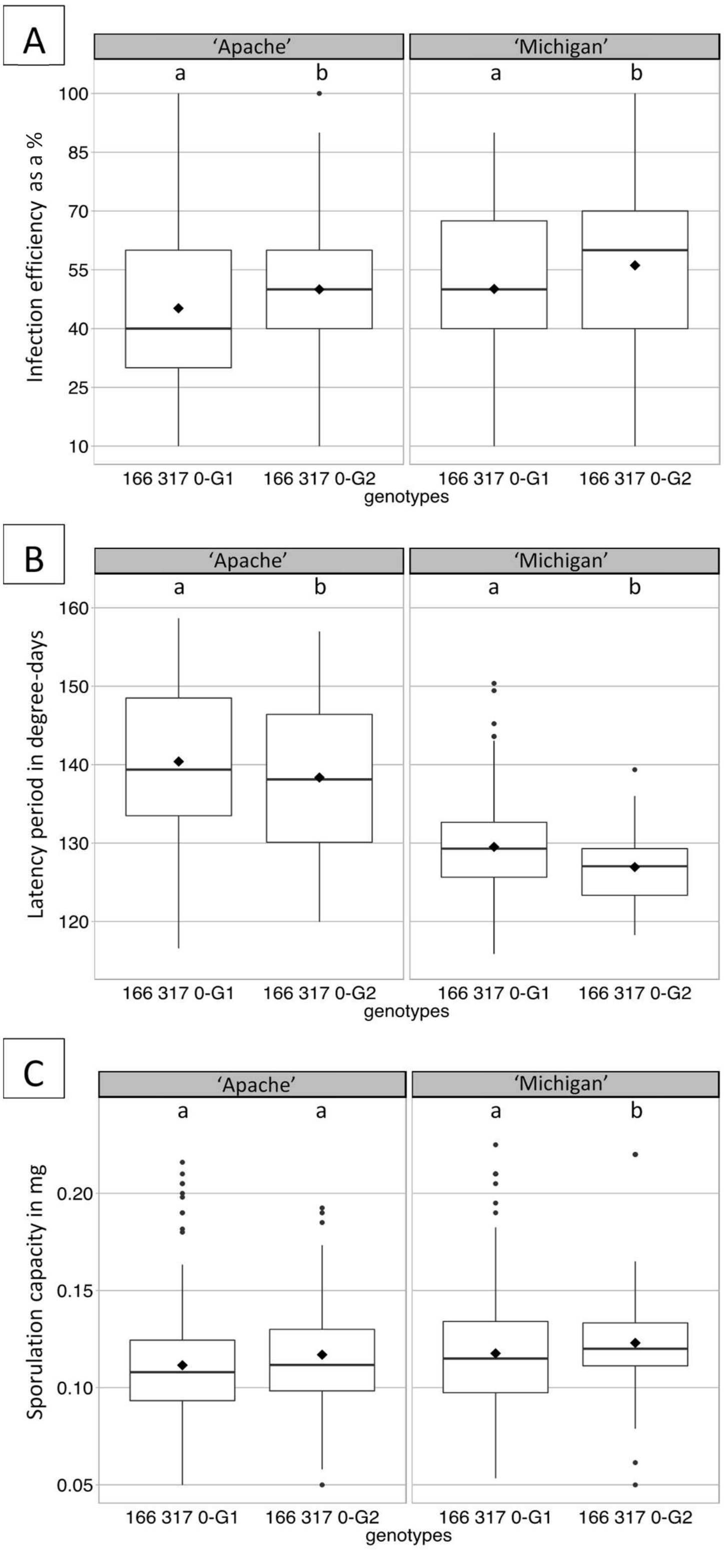


**Figure S1.** (A) Infection efficiency (IE) as a %, (B) latency period (LP) in degree-days and (C) sporulation capacity (SP) in mg of spores/uredinia for the two *Puccinia tricitina* genotypes 166 317 0-G1 and 166 317 0-G2 assessed on cultivars Apache and Michigan. Isolates were sampled from cv. Apache in 2012, 2014 and 2016. Within a box plot, the black diamond represents the mean value and the bar indicates the median. Letters indicate significant difference between genotypes, in Kruskal-Wallis tests (C) or ANOVA (A and B).

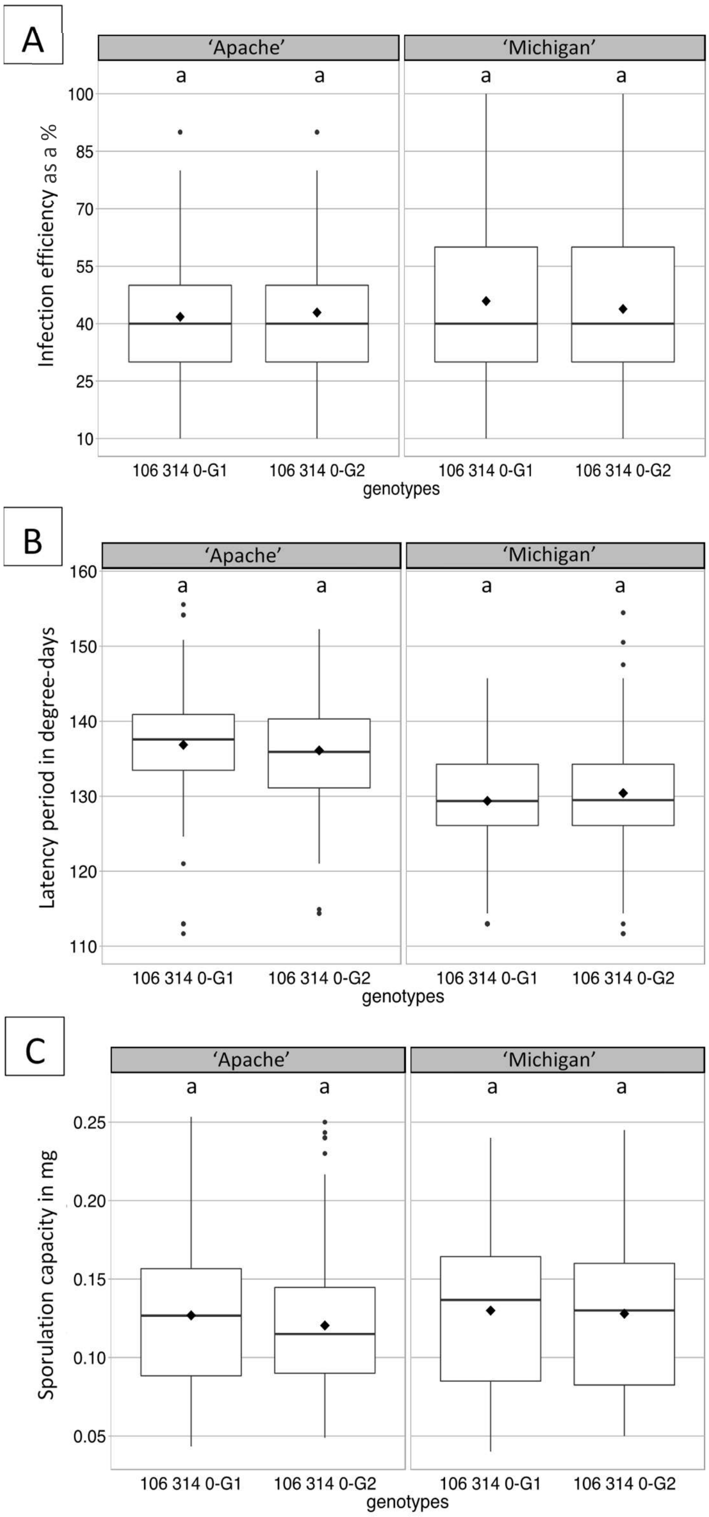


**Figure S2.** Infection efficiency as a %, latency period in degree-days and sporulation capacity in mg of spores/uredinia for *Puccinia tricitina* genotypes of pathotype 106 314 0 measured on cultivars Apache and Michigan. Isolates were sampled from cv Apache, in 2005-2006, 2012-2013 and 2015-2016. Within a box plot, the black diamond represents the mean value and the bar represents the median value. Letters indicate significant differences between genotypes in Kruskal-Wallis tests (B and C) or ANOVA (A).
